# M^6^A-modified circEZH2 Protects Endothelial Cells from Senescence and Suppresses Atherosclerosis by Stabilizing ZNF326

**DOI:** 10.64898/2026.06.05.730538

**Authors:** Xia Jing, Xue Min, Zi-yi Jin, Zhencan Shi, Yingming Li, Zhao-fu Liao, Meng-yun Cai, Xin-bin Tang, Yumin Qiu, Zi-wen Xia, Yi-tuan Xie, Yun-Fei Qu, Song-hua Wang, Lushuang Mao, Heng Li, Zhu-guo Wu, Xin-guang Liu, Jun Tao, Xing-dong Xiong

## Abstract

**BACKGROUND:** Endothelial cell senescence induces endothelial dysfunction, thereby contributing to atherosclerosis progression. Circular RNAs (circRNAs) play diverse roles in multiple physiological and pathological processes. N^6^-methyladenosine (m^6^A) is the most abundant internal RNA modification in eukaryotic RNAs and dynamically regulates RNA fate and function. However, the functions and therapeutic potential of m^6^A-modified circRNAs in endothelial cell senescence remain unknown.

**METHODS:** m6A-modified circRNAs associated with endothelial cell senescence were screened by circRNA expression and m6A-circRNA microarray profiling of endothelial cells and mouse aortic intima. circEZH2 expression was validated in endothelial cells, vascular tissues, and human atherosclerotic plaques by RT-qPCR, and RNA fluorescence *in situ* hybridization. The role of circEZH2 in endothelial senescence and atherosclerosis was assessed *in vitro* and *in vivo*. RNA pull-down, mass spectrometry, RNA immunoprecipitation, co-immunoprecipitation, ubiquitination assays, and rescue experiments were used to define the underlying mechanism.

**RESULTS:** We identified A novel m^6^A-modified circRNA, circEZH2, that was downregulated in the aged aortic intima and advanced plaques. CircEZH2 was stabilized by m6A reader IGF2BP2. Endothelial cell-specific overexpression of circEZH2 delayed senescence and suppressed atherosclerosis progression. At the cellular level, circEZH2 overexpression delayed senescence, decreased p53/p21 levels and increased angiogenic activity of endothelial cells, while circEZH2 knockdown exhibited the opposite effect. Mechanistically, circEZH2 functions as a scaffold to promote USP37-mediated deubiquitination, thereby stabilizing ZNF326. Moreover, endothelial cell-specific knockdown of ZNF326 counteracts the anti-senescent and anti-atherosclerotic effects mediated by circEZH2 overexpression.

**CONCLUSIONS:** In summary, the present study identifies circEZH2 as a novel suppressor of endothelial cell senescence, highlighting its potential as a therapeutic target for age-related atherosclerosis.

**GRAPHIC ABSTRACT:** A graphic abstract is available for this article.

Graphic abstract.
Schematic model of m^6^A-modified circEZH2 regulation in endothelial cell senescence and atherosclerosis. In young endothelial cells, IGF2BP2 is highly expressed and recognizes m^6^A-modified circEZH2, thereby maintaining its RNA stability. CircEZH2 stabilizes ZNF326 protein through USP37-mediated deubiquitination, which leads to suppression of p21 and p53, delays endothelial cell senescence, ameliorates endothelial dysfunction, and ultimately suppresses the progression of atherosclerosis.

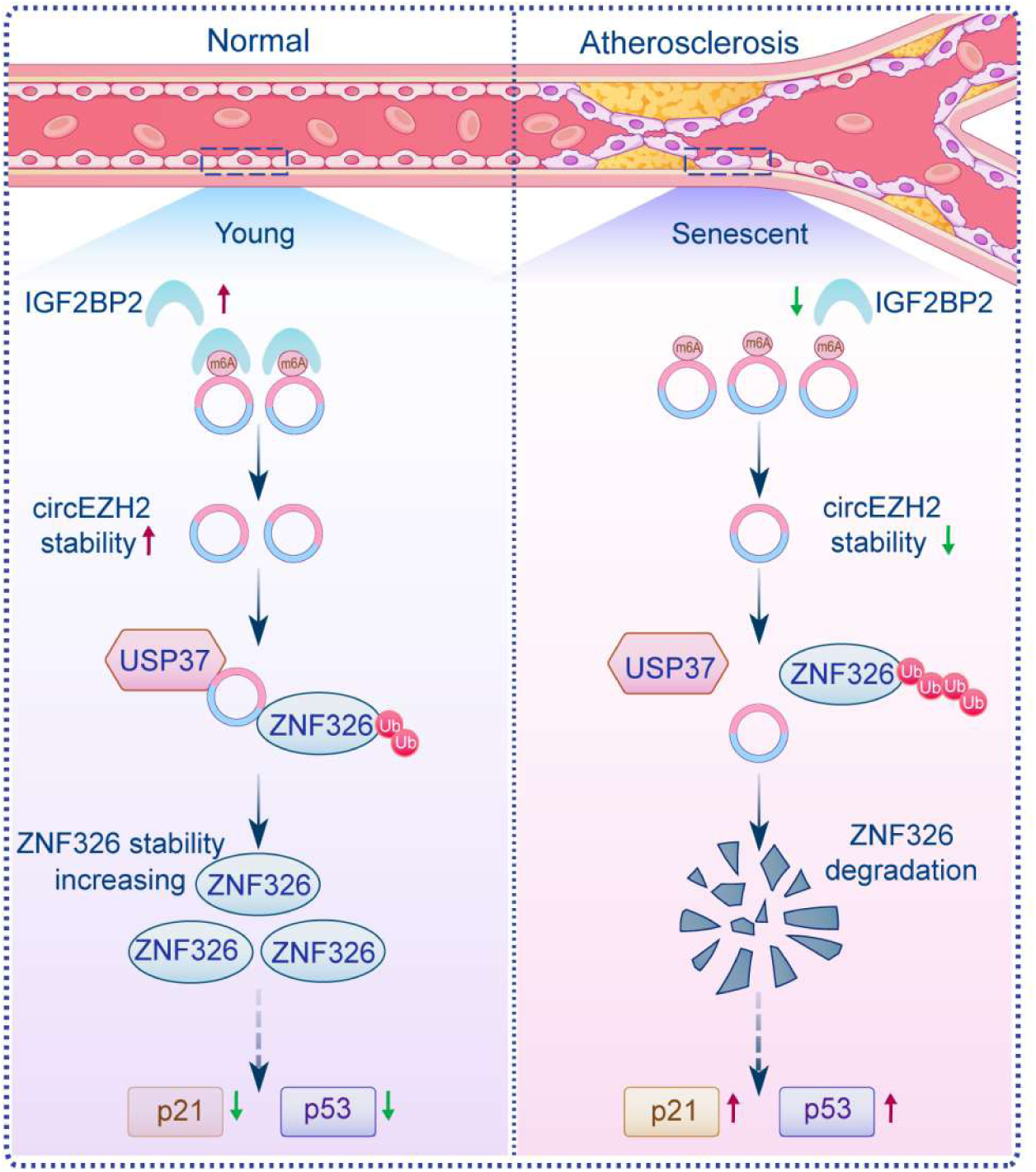

**What Are the Clinical Implications?:** This study identifies circEZH2 as a novel m6A-modified circular RNA that is reduced in the aged aortic intima and in endothelial cells within advanced atherosclerotic plaques. Endothelial circEZH2 overexpression delays endothelial cell senescence, preserves endothelial function, and suppresses atherosclerotic lesion formation, supporting an important role for circEZH2 in vascular aging-associated atherosclerosis. Mechanistically, circEZH2 acts as a scaffold to enhance USP37-mediated deubiquitination and stabilization of ZNF326, while endothelial ZNF326 knockdown counteracts the anti-senescent and anti-atherosclerotic effects of circEZH2. These findings reveal the circEZH2-ZNF326 axis as a previously unrecognized mechanism regulating endothelial senescence and atherosclerosis progression. Our work supports the potential of circEZH2-based gene therapy as a novel therapeutic approach for atherosclerosis.

## Introduction

Endothelial cells form a monolayer lining the interior surface of blood vessels and are essential for maintaining vascular homeostasis by regulating vascular tone and integrity.^1^ With aging, endothelial cells undergo morphological, functional, and gene expression alterations, including cellular enlargement, diminished proliferative capacity, and upregulation of cell cycle inhibitors such as p53 and p21.^2^ Senescent endothelial cells express increased levels of adhesion molecules and pro-inflammatory cytokines, along with reduced expression of endothelial nitric oxide synthase (eNOS).^3, 4^ These changes lead to impaired endothelium-dependent vasodilation and endothelial dysfunction, thereby contributing to the progression of atherosclerosis.^5^ Thus, exploring the mechanisms underlying endothelial cell senescence may facilitate the development of novel therapeutic strategies against atherosclerosis.

Circular RNAs (circRNAs) are covalently closed RNA molecules generated by back-splicing of pre-mRNAs, lacking both 5′ caps and 3′ poly(A) tails.^6^ They exert diverse functions by interacting with RNA-binding proteins, serving as scaffolds for protein–protein interactions, acting as microRNA sponges, regulating transcription and alternative splicing, and encoding functional peptides.^7–13^ With the advancement of high-throughput sequencing technologies and bioinformatic algorithms, an increasing number of circRNAs have been identified.^14^ Many of them are involved in a wide range of physiological and pathophysiological processes.^15–18^

N6-methyladenosine (m^6^A) is the most abundant internal modification in eukaryotic RNAs and plays essential roles in RNA metabolism, including splicing, nuclear export, stability, translation, and degradation.^19^ This modification is dynamically regulated by m^6^A “writers” (METTL3, METTL14, KIAA1429, and WTAP), “erasers” (FTO, ALKBH5 and ALKBH3), and “readers” (YTHDF and IGF2BP family proteins).^20^ Emerging evidence further indicates that m6A modification occurs not only in linear RNAs but also in circRNAs.^21^ m^6^A-modified circRNAs have been implicated in diverse biological and pathological processes, including cardiac ischemia–reperfusion injury, hypoxic stress responses and tumor progression.^22–24^ However, whether m^6^A-modified circRNAs regulate endothelial cell senescence is still unclear.

In this study, we discovered that circEZH2, which originates from exons 2 and 3 of the EZH2 gene, was significantly downregulated in the aged aortic intima and advanced plaques. Furthermore, we identify IGF2BP2-dependent m^6^A modification as a critical regulator of circEZH2 stability. Endothelial circEZH2 overexpression delayed endothelial cell senescence and suppressed atherosclerosis progression. Mechanistically, circEZH2 enhanced the effect of USP37 on ZNF326 deubiquitination and stabilization by acting as a scaffold. Taken together, we uncover a novel circRNA, circEZH2, that delays endothelial cell senescence and alleviates endothelial dysfunction, suggesting its potential as a therapeutic target for atherosclerosis.

## Materials and Methods

### Animals and human artery samples

All animal experiments were performed at the Guangdong Medical University Animal Center (Guangdong, China) under specific pathogen-free (SPF) conditions, with a 12-h light/dark cycle. Male apolipoprotein E knockout (*ApoE*^−/−^) and C57BL/6J mice were purchased from Shanghai Model Organisms Center, Inc. (Shanghai, China) and maintained in standard cages. All experimental protocols were reviewed and approved by the Animal Care and Use Committee of Guangdong Medical University, and all procedures complied with the *Guide for the Care and Use of Laboratory Animals* (NIH, 2011).

Left internal thoracic arteries (from coronary artery bypass grafting) and carotid atherosclerotic plaques (from carotid endarterectomy) were collected from patients at the Affiliated Huizhou First Hospital, Guangdong Medical University. The study was approved by the Ethics Committee of the hospital, and all procedures complied with the Declaration of Helsinki. All patients provided written informed consent prior to participation in the study.

### Mouse experiments

Recombinant AAV9 vectors carrying circEZH2 and the empty vector with the mouse endothelial-specific promoter ICAM2 (AAV9-ICAM2-circEZH2 and AAV9-ICAM2-circCon) were constructed and provided by Vigene Biosciences (Shandong, China). *ApoE⁻/⁻* mice received a tail vein injection of 1 × 10^12^ viral genomes (vg) of either AAV9-ICAM2-circEZH2 or the control vector. For the *in vivo* rescue experiments, a mixture of 5 × 10^11^ vg AAV9-ICAM2-circEZH2 and either 5 × 10^11^ vg AAV9-ICAM2-shZNF326 or AAV9-ICAM2-shRNA-Con (control) was coinjected into *ApoE⁻/⁻* mice. Mice were then fed a 12-week high-fat diet (HFD) and subsequently euthanized. The aortas and hearts with aortic roots were harvested. The degree of atherosclerosis and senescence in both the aortic root and the entire aorta was evaluated using Oil Red O staining and SA-β-gal staining.

In addition, RNA from aortic intima or media/adventitia (M/A) of mice was isolated to quantify circEZH2 expression levels. Briefly, aortas were isolated and carefully cleaned of peri-adventitial tissue. After flushing the lumen with PBS, TRIzol was flushed into the lumen using a 29-gauge insulin syringe, and the eluate containing the aortic intima was harvested into a microcentrifuge tube. The remaining arterial media and adventitia were also collected in TRIzol for subsequent RNA extraction.

### Cell culture

Primary human umbilical vein endothelial cells (HUVECs), human aortic endothelial cells (HAECs) and human coronary artery endothelial cells (HCAECs) were obtained from ScienCell Research Laboratories (USA). The cells were cultured in Endothelial Cell Medium (ECM; ScienCell) supplemented with 5% fetal bovine serum (FBS), 1% endothelial cell growth supplement (ECGS), and 1% penicillin–streptomycin solution at 37 °C in a humidified atmosphere containing 5% CO₂. Replicative senescence model was induced in endothelial cells by serial passaging. HUVECs at PDL < 8 and at PDL > 29 were defined as proliferating and senescent cells, respectively. HAECs at PDL < 3 and PDL > 13 were defined as proliferating and senescent cells, respectively. HCAECs at PDL < 4 were defined as proliferating cells, and PDL > 20 as senescent cells. For H_2_O_2_-induced senescence, proliferating endothelial cells were exposed to 100 μmol/L H_2_O_2_ for 1 h and then cultured in complete medium.

### RNA extraction and RT-qPCR

Total RNA was extracted from the samples using TRIzol reagent (Invitrogen, USA), and reverse transcribed into cDNA with HiScript III RT SuperMix for qPCR (+gDNA wiper) (Vazyme, China). qPCR was performed on an ABI QuantStudio 3 Real-Time PCR System (Thermo Fisher Scientific, USA) using Taq Pro Universal SYBR qPCR Master Mix (Vazyme, China) to quantify circRNA and mRNA levels. β-actin served as the internal reference. Relative expression was calculated with the 2^−ΔΔCt^ method. Primer sequences are provided in Supplementary Table S3.

### circRNA microarray analysis

Total RNA extracted from the aortic intima of young and aged mice was treated with RNase R (Epicentre, USA) to remove linear RNAs and enrich circRNAs. The enriched circRNAs were amplified and transcribed into fluorescent complementary RNAs (cRNAs) using random primers, following the manufacturer’s protocol for the Arraystar Super RNA Labeling Kit (Arraystar Inc., USA). Labeled cRNAs were then hybridized to the Arraystar Mouse circRNA Microarray V2 (8×15K, Arraystar) and incubated at 65 °C for 17 h in an Agilent Hybridization Oven. After washing, the arrays were scanned on an Agilent G2505C Scanner. Raw data were extracted with Agilent Feature Extraction software (v11.0.1.1). Subsequent data normalization and analysis were performed with the R package limma.

### m^6^A-circRNA epitranscriptomic microarray

For m^6^A-circRNA epitranscriptomic microarray analysis, total RNA extracted from HUVECs and mouse aortic intimal tissues was immunoprecipitated with an anti-m^6^A antibody. The immunoprecipitated RNA and the corresponding supernatant RNA were collected as the immunoprecipitated (IP) and supernatant (Sup) fractions, respectively, followed by RNase R digestion to enrich circRNAs. The enriched IP and Sup RNAs were then labeled with Cy5 and Cy3, respectively, and hybridized onto the Arraystar m^6^A-circRNA Epitranscriptomic Microarray according to the manufacturer’s instructions. Raw data were extracted and normalized to identify m^6^A-modified circRNAs.

### RNA immunoprecipitation (RIP) and m^6^A RIP (MeRIP)

RNA immunoprecipitation was performed using the Magna RIP RNA-Binding Protein Immunoprecipitation Kit (Millipore, USA) following the manufacturer’s instructions. Magnetic beads conjugated with anti-IGF2BP2 (Proteintech, China), anti-YTHDF2 (Cell Signaling Technology, USA), anti-USP37 (Proteintech, China) or control IgG (Proteintech, China) antibodies were incubated with cell lysates overnight at 4 °C on a rotator. The co-immunoprecipitated RNA and protein were subsequently analyzed by RT-qPCR and Western blot, respectively.

MeRIP assays were performed using the Magna MeRIP m^6^A Kit (Millipore, USA) according to the manufacturer’s protocol. Briefly, total RNA was fragmented and incubated with magnetic beads conjugated with anti-m^6^A antibody (Abcam, UK) or control IgG at 4°C overnight. The immunoprecipitated methylated RNA was then eluted, purified, and analyzed by RT-qPCR. The relative enrichment of circEZH2 was normalized to the input.

### RNase R and actinomycin D (Act D) treatment

Total RNA (5 μg) from HUVECs was incubated for 15 min at 37 °C with or without RNase R (Epicentre, USA), followed by RT-qPCR analysis of circEZH2 and EZH2 mRNA levels. For Act D treatment, Act D (Sigma-Aldrich, USA) was added to the culture medium at a final concentration of 10 μg/mL. Cells were harvested at 0, 6, 12, 18, 24 h after treatment, and total RNA was extracted for RT-qPCR analysis.

### RNA fluorescence in situ hybridization (RNA-FISH)

RNA-FISH was performed using a method based on padlock probes (PLPs) and rolling circle amplification (RCA) to localize circEZH2 in cultured cells and tissue sections. Briefly, samples were fixed in 4% paraformaldehyde for 30 min. Padlock probes specific to circEZH2 were designed to target its back-splice junction sequence (Table S3). Following 4 h hybridization with the phosphorylated padlock probes at 37 °C, ligation was carried out at 37 °C for 1 h using a mix containing 1× SplintR buffer (NEB, USA), 2.5 U/μL SplintR Ligase (NEB, USA), 50% glycerol (Sigma-Aldrich, USA), and 0.2 μg/μL BSA (Solarbio, China), resulting in the formation of circular DNA molecules. RCA primers were then hybridized to the circular DNA templates at 37 °C for 30 min. Rolling circle amplification was subsequently conducted with phi29 DNA polymerase (Thermo Fisher Scientific, USA) at 37 °C overnight, generating RCA products composed of numerous repeated complementary sequences of the original DNA circles. Finally, fluorophore-conjugated detection probes were hybridized to the RCA products for 30 min at room temperature. Nuclei were counterstained with DAPI, and fluorescence signals were visualized using a confocal laser scanning microscope (Leica TCS SP8; Leica Microsystems, Germany).

### Transfection and lentivirus infection

Two independent siRNAs against the back-splice junction of circEZH2 and two siRNAs against IGF2BP2 were synthesized by RiboBio (Guangzhou, China). Transfection was carried out using Lipofectamine RNAiMAX reagent (Life Technologies, USA) following the manufacturer’s protocol. The silencing efficiency was validated by RT-qPCR.

For circEZH2 overexpression, lentiviral vectors were purchased from GeneChem Co., Ltd. (Shanghai, China). HUVECs were infected with Lv-circEZH2 or Lv-circCon (lentiviral control) at a multiplicity of infection (MOI) of 10. After incubation for 12-24 h, the medium was replaced with fresh culture medium. Infected cells were selected with puromycin (2 μg/mL) for 5-7 d to establish stable expression.

### Senescence-associated β-galactosidase (SA-β-gal) staining

Senescence-associated β-galactosidase (SA-β-gal) staining was performed using the Senescence β-Galactosidase Staining Kit (Beyotime, China) following the manufacturer’s instructions. Briefly, after washing twice with PBS, HUVECs or mouse aortic root sections were fixed in the fixative solution for 15 min at room temperature. The samples were then washed and incubated with freshly prepared SA-β-gal staining solution overnight at 37°C in a dry, CO₂-free incubator. Stained cells or sections were imaged under a light microscope. For whole aortas, tissues were harvested from sacrificed mice and perfused with ice-cold PBS, followed by staining according to the SA-β-gal kit protocol.

### BrdU incorporation assay

Cell proliferation was assessed using a BrdU (5-bromo-2-deoxyuridine) incorporation assay. HUVECs were incubated with 40 μM BrdU (Sigma-Aldrich, USA) for 1 h at 37 °C in culture medium. After incubation, cells were fixed with 4% paraformaldehyde for 30 min at room temperature and then permeabilized with 0.05% trypsin for 7 min. Subsequently, the cells were blocked with 3% bovine serum albumin (BSA) for 1 h. Incorporated BrdU was detected using a mouse anti-BrdU monoclonal antibody (Sigma-Aldrich, USA), followed by incubation with an Alexa Fluor 488-conjugated secondary antibody (Cell Signaling Technology, USA). Nuclei were counterstained with DAPI (Invitrogen, USA), and images were captured at 100× magnification using the EVOS FL Auto Imaging System (Thermo Fisher Scientific, USA). The proliferation rate was quantified as the percentage of BrdU-positive nuclei relative to the total number of nuclei.

### Tube formation assay

The tube formation ability of endothelial cells was assessed through a Matrigel-based in vitro angiogenesis assay. Briefly, Matrigel matrix (BD Biosciences, USA) was thawed on ice and added (150 μL/well) into pre-cooled 48-well plates, followed by incubation at 37 °C for 1 h to allow gel polymerization. HUVECs were seeded onto the Matrigel-coated wells at a density of 3 × 10^4^ cells per well. After incubation for 4-6 h at 37 °C, tube-like structures were imaged at 100 × magnification using an Eclipse TS100 inverted microscope (Nikon, Japan). Tube formation was quantified by measuring tube length with ImageJ software.

### Atherosclerotic lesion analysis

Mice were euthanized, and the hearts and aortas were perfused with PBS via the left ventricle. Hearts were harvested, embedded in OCT compound (Sakura Finetek, Japan), and cut into 8 μm thickness with a cryostat (Thermo Fisher Scientific, USA) to obtain aortic root cross-sections. Sections were stained with Oil Red O and counterstained with Mayer’s hematoxylin. Atherosclerotic lesion areas were quantified using ImageJ software. The percentage of plaque area was calculated as the Oil Red O-positive (Oil Red O^+^) area divided by the total aortic root area.

For *en face* analysis, the entire aorta from the ascending arch to the iliac bifurcation was carefully dissected and cleared of surrounding adipose tissue. After longitudinally opening the vessel, samples were fixed in 10% neutralized formalin, rinsed with 60% isopropanol, and stained with freshly prepared Oil Red O solution for 10 min at room temperature. Stained aortas were briefly washed in 60% isopropanol and PBS, then placed with the intimal surface facing upward for photography with a digital camera. Plaque area was measured using ImageJ, and the percentage of plaque area was calculated as the ratio of Oil Red O^+^ area to total aortic area.

### Immunofluorescence staining

Frozen tissue sections were fixed in 4% paraformaldehyde for 15 min at room temperature, followed by three washes with PBS. Samples were then permeabilized using 0.3% Triton X-100 in PBS (PBST) for 10-15 min and subsequently blocked with 10% goat serum in PBST for 1 h at room temperature. Primary antibodies, including anti-CD31 (1:150, BioLegend, USA), anti-IGF2BP2 (1:100, Proteintech, China), anti-p53 (1:200, Proteintech, China), anti-p21 (1:100, Cell Signaling Technology, USA), anti-ZNF326 (1:100, Proteintech, China), anti-VCAM1 (vascular cell adhesion molecule 1, 1:400, Cell Signaling Technology, USA), and anti-ICAM1 (intercellular adhesion molecule 1, 1:200, Santa Cruz, USA) were diluted in 0.3% PBST and incubated with the sections at 4 °C overnight in a humidified chamber. After washing with PBS, sections were incubated with fluorophore-labelled secondary antibodies in PBST for 1 h at room temperature in the dark. Nuclei were counterstained with DAPI for 10 minutes. After a final PBS wash, the sections were mounted using anti-fade mounting medium. Images were acquired using a confocal laser scanning microscope (Zeiss LSM980).

### CircRNA pull-down assay and mass spectrometry

The circRNA pull-down assay was performed using the MS2-tagged system to identify RNA-binding proteins (RBPs) interacting with circEZH2. Briefly, expression vectors carrying circEZH2-MS2 and MS2-CP-Flag were constructed (Geneseed, China) and co-transfected into HUVECs. Upon intracellular expression, MS2-CP specifically binds to the MS2-tagged circEZH2, thereby forming a stable MS2-CP-MS2-circEZH2 complex. Subsequently, cell lysates were prepared in lysis buffer supplemented with RNase and protease inhibitors. The MS2-CP-MS2-circEZH2 complex was immunoprecipitated using anti-Flag antibody pre-conjugated to Protein A/G magnetic beads. After incubation at 4 °C for 1-2 h, the beads were thoroughly washed with washing buffer to remove non-specific interactions. The bound protein complexes were then eluted and subjected to mass spectrometry analysis (Q Exactive, Thermo Scientific, USA).

### Cycloheximide (CHX) treatment

HUVECs were transfected with si-NC or si-circEZH2 and then treated with 10 μg/mL cycloheximide (CHX, Sigma, USA), a protein synthesis inhibitor, for 0, 5, 10, or 20 h. Proteins were subsequently extracted from the treated cells, and ZNF326 expression levels were analyzed by Western blot.

### Ubiquitination assay

Cells were transfected with si-NC or si-circEZH2, with or without co-transfection of HA-ZNF326 and His-tagged ubiquitin (His-Ub) expression vectors. Following transfection, cells were treated with 10 μM MG132 (MCE, USA) for 8 h, after which total proteins were extracted for immunoprecipitation with anti-ZNF326 (Santa Cruz, USA) or anti-HA (Proteintech, China) antibodies. The ubiquitination of ZNF326 was subsequently detected by Western blot using anti-ubiquitin (Cell Signaling Technology, USA) or anti-His (Proteintech, China) antibodies.

### Co-immunoprecipitation (Co-IP)

Cells were harvested and lysed in IP lysis buffer with protease inhibitors on ice for 20 min, followed by centrifugation at 16,000 × g for 10 min at 4 °C. The supernatant was incubated with the indicated antibodies plus Protein A/G magnetic beads (Thermo Fisher Scientific, USA) overnight at 4 °C on a rotator. Beads were collected using a magnetic stand, washed with wash buffer, and then boiled in 1× SDS-loading buffer at 100 °C for 10 min. The immunoprecipitated proteins were subsequently analyzed by Western blot.

### Western blot analysis

Total proteins from cells and tissues were extracted using RIPA lysis buffer (Beyotime, China) supplemented with protease inhibitors. Equal amounts of protein were separated on 12.5% SDS–PAGE and transferred to PVDF membranes (Millipore, USA). The membranes were blocked with 5% nonfat milk in TBST for 1 h at room temperature, then incubated overnight at 4 °C with primary antibodies against IGF2BP2 (Proteintech, China), p53 (Cell Signaling Technology, USA), p21 (Cell Signaling Technology, USA), GAPDH (Proteintech, China) and ZNF326 (Proteintech, China) diluted according to the manufacturers’ instructions. After washing, membranes were incubated with the appropriate HRP-conjugated secondary antibodies (Beyotime, China) for 1-2 h at room temperature. Protein bands were visualized with Immobilon Western Chemiluminescent HRP Substrate (Millipore, USA) and quantified in ImageJ software.

### Statistical analysis

All statistical analyses were performed using GraphPad Prism 9.0 (GraphPad Software, USA). Data are presented as the mean ± standard deviation (SD). Normality and homogeneity of variance were assessed prior to data analysis. For comparisons between two groups, a two-tailed unpaired Student’s *t*-test was used. For comparisons among more than two groups, one-way ANOVA or Welch’s ANOVA was applied depending on the homogeneity of variances. A *p*-value < 0.05 was considered statistically significant.

## Results

### circEZH2 is downregulated in the aged aortic intima and advanced plaques

To elucidate the role of m^6^A-modified circRNAs in endothelial cell senescence, circRNA microarray analysis was conducted on aortic intima samples from young and aged mice, and m^6^A-circRNA epitranscriptomic microarray analysis was performed in HUVECs and mouse aortic intimal tissues. By integrating our previously published RNA-seq dataset of young and senescent endothelial cells (GEO: GSE151475) with circRNA microarray data from young and aged aortic intima, we identified 458 conserved circRNAs that were commonly differentially expressed in both endothelial cells and aortic intimal tissues (Figure S1-2). Further intersection of these 458 circRNAs with m^6^A-modified circRNAs detected in HUVECs and mouse aortic intima yielded 211 circRNAs (Figure 1A). Among them, circEZH2, also known as hsa_circ_0006357/mmu_circRNA_0001471, was selected for further investigation because it was m^6^A-modified and significantly downregulated in both senescent endothelial cells and aged aortic intima (Table S1). Homology analysis revealed that circEZH2 shares 96% sequence identity between humans and mice (Figure S3). circEZH2 is derived from exons 2–3 of the EZH2 gene with a length of 253 nt.^25^ Its circular form was confirmed by Sanger sequencing of the back-splice junction site, and its stability was further validated by exonuclease RNase R and Actinomycin D treatment in endothelial cells (Figure S4). RNA-FISH revealed that circEZH2 expression was downregulated in the intima of aged human arteries and aged human vascular tissues (Figure 1B-C and Figure S5). In *in vitro* cultured endothelial cells undergoing replicative senescence and H_2_O_2_-induced premature senescence, circEZH2 expression was also significantly reduced (Figure 1D-E and Figure S6-7). Consistently, RT-qPCR confirmed that circEZH2 was significantly reduced in the aortic intima and vascular tissues of aged mice (Figure 1F and Figure S8). Additionally, circEZH2 expression was decreased in the aortas of *ApoE^-/-^* mice after a high-fat diet (HFD) for 12 weeks (Figure S9). In human advanced plaques, circEZH2 expression in endothelial cells was also downregulated compared with early plaques (Figure 1G-H). Collectively, these data suggest a potential role of circEZH2 in regulating endothelial cell senescence and atherosclerosis.

**Figure 1.**
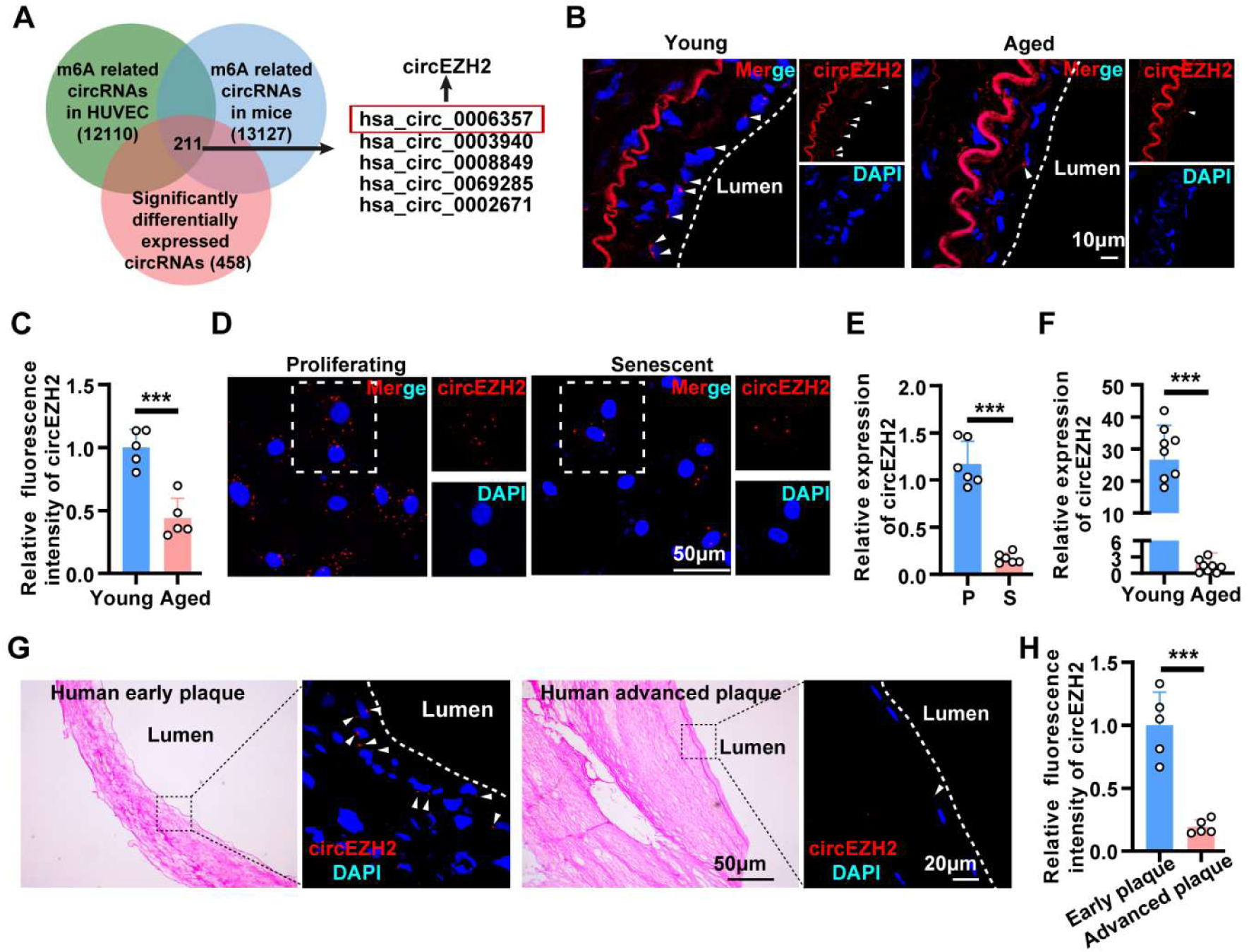
circEZH2 is downregulated in the aged aortic intima and advanced plaques. **A**,Venn diagram showing the overlap among m^6^A-modified circRNAs detected in HUVECs (green), m^6^A-modified circRNAs detected in mouse aortic intima (blue), and 458 conserved circRNAs differentially expressed in both proliferating vs. senescent endothelial cells and young vs. aged aortic intima (pink). circEZH2 (hsa_circ_0006357) was identified from the overlapping candidates and selected for further investigation. **B and C**, RNA fluorescence in situ hybridization (RNA-FISH) was performed with Cy3-labeled circEZH2 probes (red) to detect circEZH2 in young and aged human arteries. **D**, circEZH2 expression was further examined by RNA-FISH in proliferating (P) and senescent (S) HUVECs. **E**, RT-qPCR assays were conducted to detect the amount of circEZH2 in proliferating and senescent HUVECs. **F**, RT-qPCR analysis of circEZH2 levels in the intima of young (n = 8, 2 months old) and aged (n = 8, 26-28 months old) C57BL/6J mice. **G and H**, RNA-FISH and quantification analysis were performed to detect circEZH2 expression (red) in early and advanced human carotid plaques. Data are presented as means ± SD, Student’s *t*-test, \*\*\**P* < 0.001.

### circEZH2 is stabilized by m6A reader IGF2BP2

To investigate whether m^6^A modification might contribute to the altered expression of circEZH2 during endothelial cell senescence, potential m^6^A modification sites in circEZH2 were predicted using circPRIMER, revealing seven putative m^6^A sites (Figure 2A). MeRIP-qPCR assays revealed increased m^6^A enrichment of circEZH2 in senescent endothelial cells and aged aortic intima (Figure 2B and C). We then investigated whether m^6^A reader proteins were involved in circEZH2 regulation. RIP assays demonstrated that circEZH2 directly interacted with m^6^A reader IGF2BP2 but not YTHDF2 (Figure 2D-E). Functionally, IGF2BP2 knockdown reduced circEZH2 expression and accelerated circEZH2 degradation after actinomycin D treatment, suggesting that IGF2BP2 stabilizes m^6^A-modified circEZH2 in endothelial cells (Figure 2F-G). Furthermore, IGF2BP2 protein expression was decreased in senescent endothelial cells and was also reduced in the intimal layer of aged human arteries (Figure 2H-I). Together, these results indicate that circEZH2 undergoes m^6^A modification and is stabilized by the m^6^A reader IGF2BP2.

**Figure 2.**
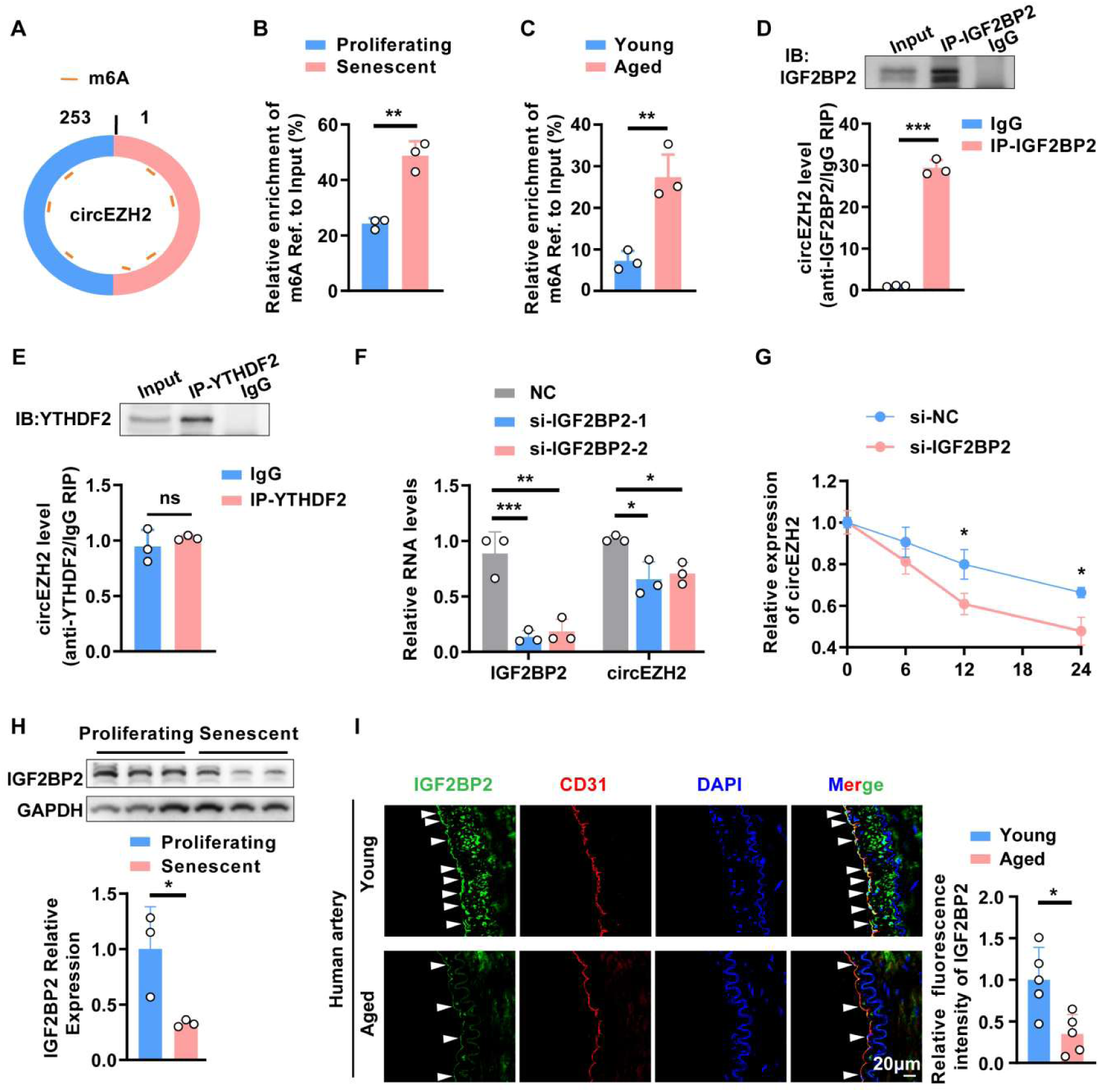
circEZH2 is an m^6^A-modified circRNA that interacts with IGF2BP2 in endothelial cells. **A**, Prediction of putative m^6^A modification sites in circEZH2 using circPRIMER. **B and C**, MeRIP-qPCR assays showing the relative m^6^A enrichment of circEZH2 in proliferating vs. senescent HUVECs (B) and in young vs. aged mouse aortic intima (C). **D and E**, RIP assays demonstrating the interaction of circEZH2 with IGF2BP2 (D) but not YTHDF2 (E). Representative immunoblots (top) and quantification of circEZH2 enrichment relative to IgG control (bottom) are shown. **F and G**, Knockdown of IGF2BP2 by two independent siRNAs (si-IGF2BP2-1 and si-IGF2BP2-2) reduced circEZH2 expression (F) and accelerated circEZH2 degradation after Actinomycin D treatment (G). **H**, Western blot analysis of IGF2BP2 protein levels in proliferating (P) and senescent (S) HUVECs. **I**, Representative immunofluorescence staining images of IGF2BP2 (green), CD31 (red) and nuclei (blue) in the intimal layer of young and aged human arteries. Data are presented as means ± SD, Student’s *t*-test; \**P* < 0.05, \*\**P* < 0.01, \*\*\**P* < 0.001.

### circEZH2 suppresses cellular senescence in endothelial cells

To evaluate the biological function of circEZH2 in endothelial cell senescence, we specifically silenced circEZH2 with small interfering RNAs (si-circEZH2) targeting the back-splice sequence and overexpressed circEZH2 with a lentivirus containing the circEZH2 coding sequence (Lv-circEZH2) in endothelial cells. We found that circEZH2 knockdown suppressed proliferation and triggered an enlarged morphology accompanied by increased SA-β-gal activity in endothelial cells, while circEZH2 overexpression exhibited the opposite effect (Figure 3A-F). Western blot analysis further indicated that the expression levels of p53 and p21, key regulators of cellular senescence, were elevated in endothelial cells with circEZH2 silencing (Figure 3G). Moreover, overexpression of circEZH2 in endothelial cells led to decreased p53 and p21 expression levels (Figure 3H). Notably, senescent endothelial cells also have impaired angiogenic capacity.^26^ As illustrated in Figure 3I and 3J, decreased circEZH2 expression impaired the tube formation ability of endothelial cells, whereas overexpression of circEZH2 enhanced tube formation ability. Taken together, our results showed that circEZH2 could play a suppressive role in endothelial cell senescence.

**Figure 3.**
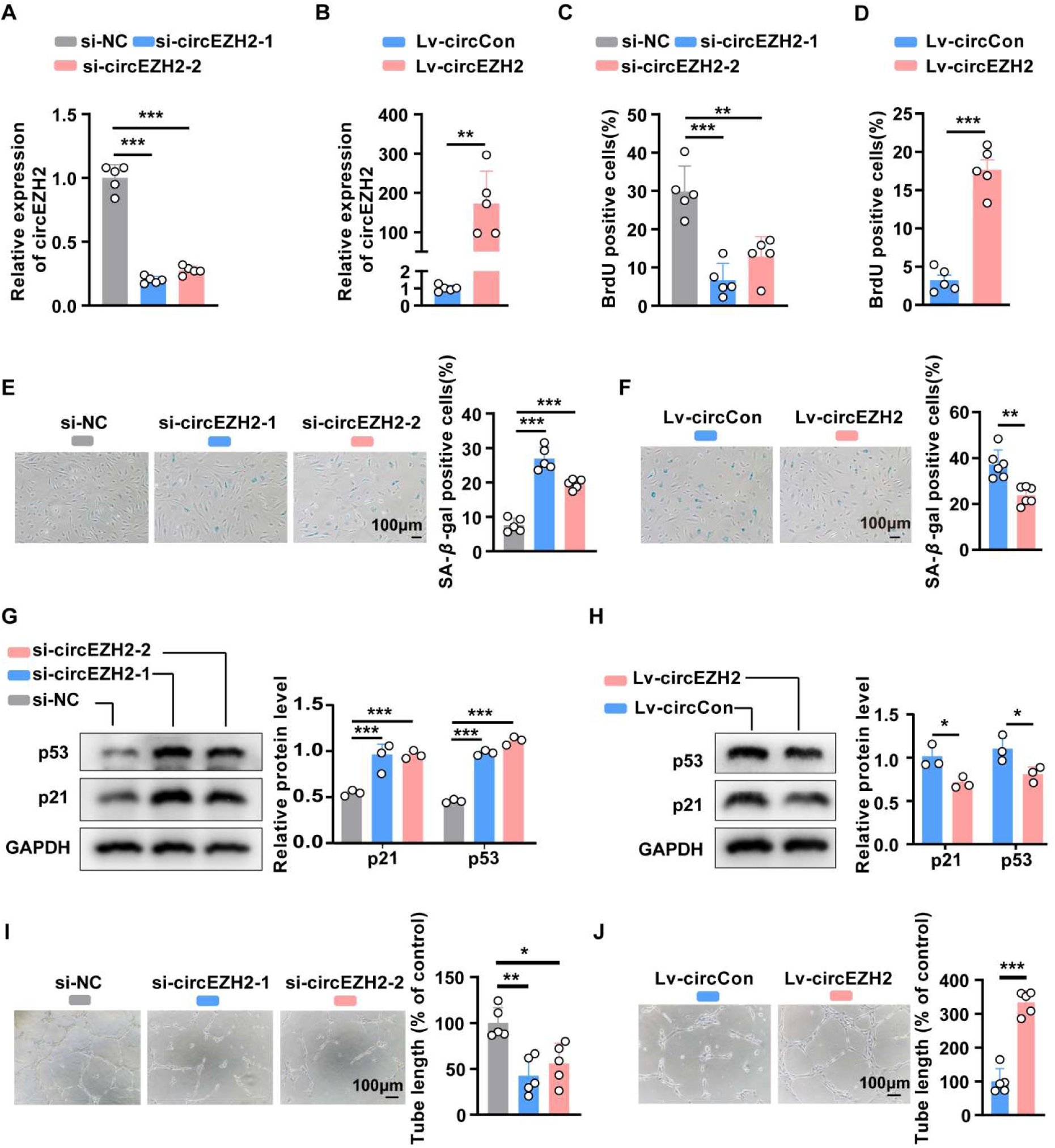
CircEZH2 suppresses cellular senescence in endothelial cells. **A**, RT-qPCR analysis of the levels of circEZH2 in young HUVECs after transfection of negative control (si-NC) or si-circEZH2-1/2. **B**, RT-qPCR analysis of the levels of circEZH2 in senescent HUVECs after infection with lentiviruses expressing circCon or circEZH2. **C, E, and I**, BrdU assays, SA-β-gal staining, and tube formation in young HUVECs after transfection with si-NC or si-circEZH2-1/2. **D, F, and J**, BrdU assays, SA-β-gal staining, and tube formation in senescent HUVECs after infection with lentiviruses expressing circCon or circEZH2. **G**, Western blot analysis of p53/p21 in young HUVECs after transfection of si-NC or si-circEZH2-1/2. **H**, Western blot analysis of p53/p21 in senescent HUVECs after infection with lentiviruses expressing circCon or circEZH2. Data are presented as means ± SD, Student’s *t*-test in B, D, F, H and J, one-way ANOVA in A, C, E, G and I, \**P* < 0.05, \*\**P* < 0.01, \*\*\**P* < 0.001.

### circEZH2 overexpression in vascular endothelium suppresses endothelial cell senescence and atherosclerosis *in vivo*

The downregulation of circEZH2 expression in the endothelium of aged aorta and advanced plaques prompted us to investigate its role in endothelial cell senescence and atherosclerosis *in vivo*. We then conducted endothelium-specific overexpression of circEZH2 in *ApoE*^-/-^ mice using AAV9, with AAV9-circCon carrying an empty vector as a control. As shown in Figure 4B, the expression of circEZH2 in the aortic intima of mice injected with AAV9-circEZH2 was 2.8-fold higher than that in mice injected with AAV9-circCon. Reduced SA-β-gal-positive areas were observed in the aortas and aortic root cross-sections of mice injected with AAV9-circEZH2, compared to AAV9-circCon (Figure 4C-D). Consistently, immunofluorescence staining demonstrated that AAV9-circEZH2 treatment led to reduced expression of p53 and p21 in the intima (Figure 4E-F). Together, these results suggested that circEZH2 could inhibit endothelial cell senescence *in vivo*.

**Figure 4.**
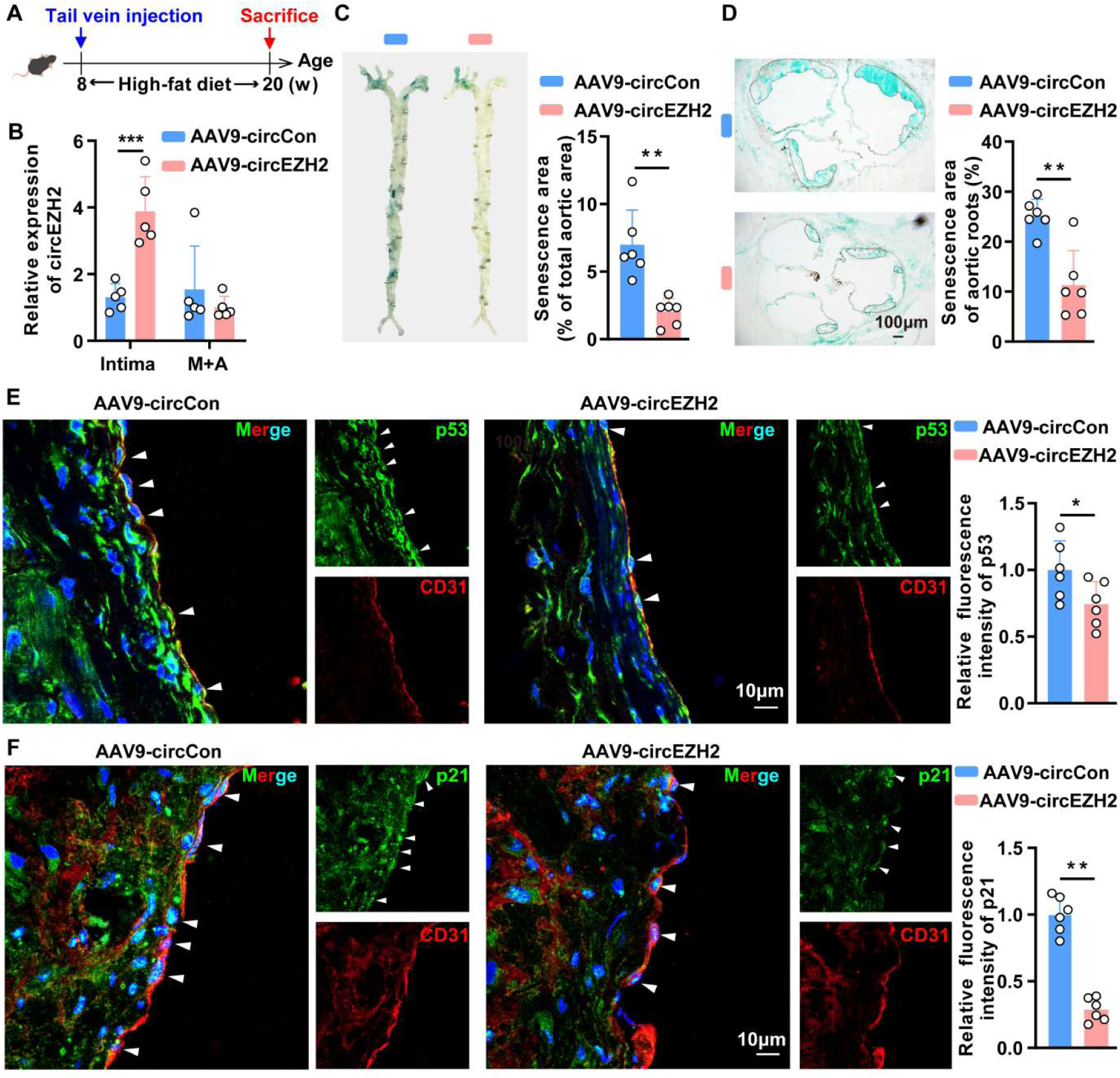
Endothelial-specific circEZH2 overexpression delays endothelial cell senescence. **A**, Schematic diagram of the time nodes of tail vein injection of AAV9-circCon and AAV9-circEZH2. **B**, RT-qPCR was used to assess circEZH2 overexpression efficiency. **C and D**, SA-β-gal staining of whole aorta and aortic roots of *ApoE^-/-^* mice following injection of AAV9-circCon and AAV9-circEZH2. **E and F**, Representative immunofluorescence staining images of CD31 (red), p53/p21 (green), and nuclei (blue) in aortic roots of *ApoE^-/-^* mice injected with AAV9-circCon and AAV9-circEZH2 (n=6 mice per group). Data are presented as means ± SD, Student’s *t*-test, \**P* < 0.05, \*\**P* < 0.01.

Endothelial cell senescence leads to endothelial dysfunction, thereby contributing to the progression of atherosclerosis.^27^ In this study, *ApoE*^-/-^ mice with endothelial cell-specific circEZH2 overexpression exhibited significantly reduced atherosclerotic plaques in the aortas and aortic root cross-sections compared to control mice (Figure 5A-C). Furthermore, AAV9-circEZH2 treatment increased eNOS levels and reduced VCAM1 and ICAM1 levels in the endothelium compared to AAV9-circCon (Figure 5D-F). Collectively, these findings indicate that circEZH2 could inhibit endothelial cell senescence, protect against endothelial dysfunction and atherosclerosis.

**Figure 5.**
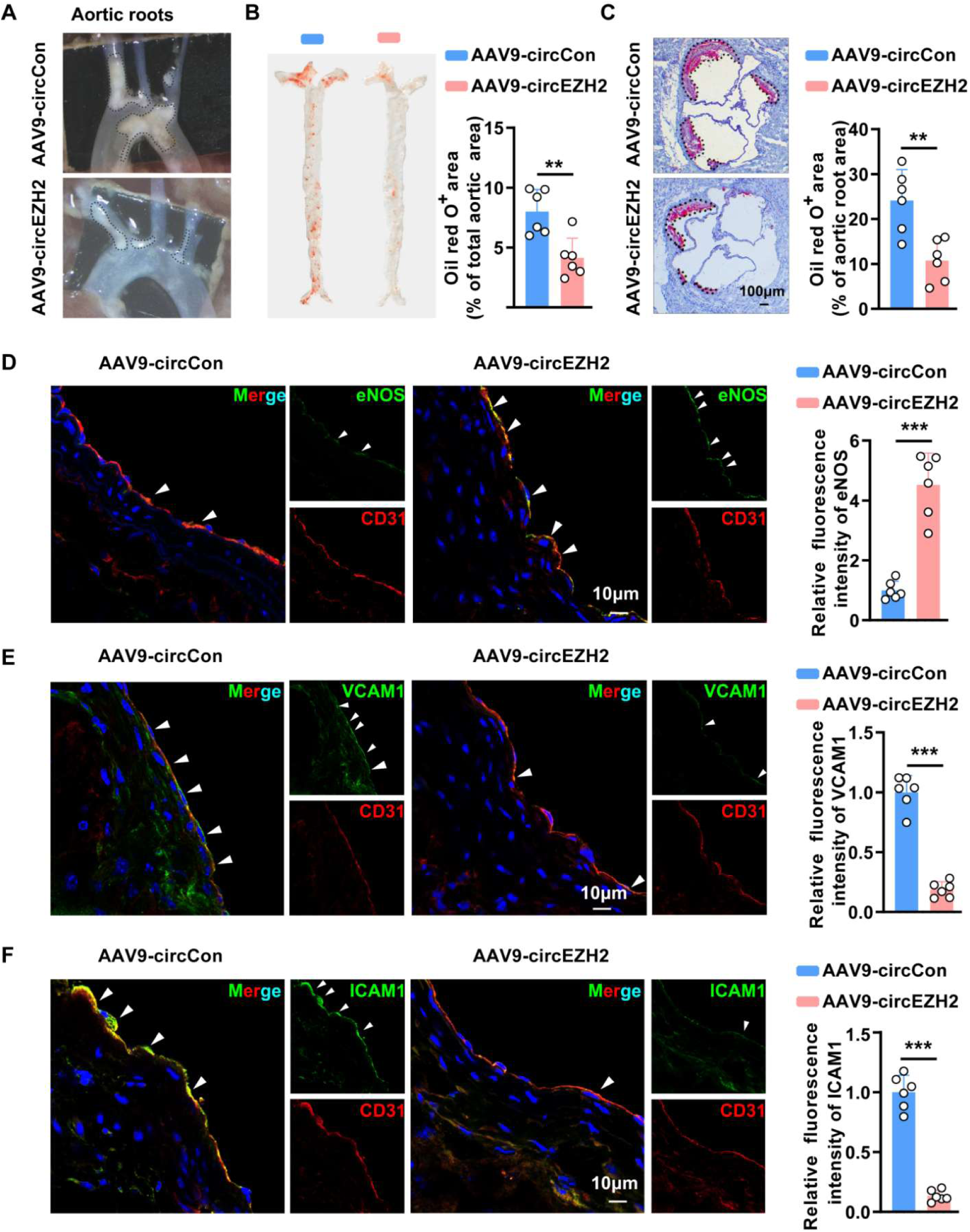
Endothelial cell-specific overexpression of circEZH2 suppresses atherosclerosis. **A**, Plaques in the aortic arches of *ApoE^-/-^* mice following injection of AAV9-circCon and AAV9-circEZH2 (n = 6 mice per group). **B and C**, Oil Red O staining of whole aorta and aortic roots of *ApoE^-/-^* mice following injection of AAV9-circCon and AAV9-circEZH2 (n = 6 mice per group). **D-F**, Representative immunofluorescence staining images of CD31 (red), eNOS/VCAM1/ICAM1 (green), and nuclei (blue) in aortic roots of *ApoE^-/-^* mice injected with AAV9-circCon and AAV9-circEZH2 (n = 6 mice per group). Data are presented as means ± SD, Student’s *t*-test, \*\**P* < 0.01, \*\*\**P* < 0.001.

### ZNF326 acts as a partner of circEZH2 to delay endothelial cell senescence

To elucidate the molecular mechanisms by which circEZH2 regulates endothelial cell senescence, we examined its effects on the expression of its host gene, EZH2. We found that altering circEZH2 expression by either overexpression or silencing affected only the circular transcript, with no impact on the linear form of EZH2 (Figure 3A-B and S11). Considering that circRNAs commonly function through protein binding, we performed an MS2-CP-Flag circRNA pull-down assay coupled with mass spectrometry (MS) to identify the protein partners of circEZH2 (Figure 6A). MS analysis identified 15 proteins enriched in the circEZH2-MS2+MS2-CP pull-down samples, with ZNF326 ranked the highest according to the peptide identification results (Figure S12 and Table S2). Next, an RNA immunoprecipitation (RIP) assay and RT-qPCR analysis confirmed the interaction between circEZH2 and ZNF326 in HUVECs (Figure 6B). ZNF326 is a multifunctional RNA/DNA-binding protein involved in RNA splicing, DNA damage response, and cellular stress pathways.^28–30^ Its dysregulation has been associated with cancer progression, including glioblastoma and breast cancer.^28, 29^ In this study, ZNF326 expression was significantly downregulated in the aortic intima of aged mice, senescent endothelial cells, and whole aortas from aged mice (Figure 6C-F and Figure S13). To directly observe the effect of ZNF326 on endothelial cell senescence, two siRNAs were designed to target ZNF326. The deficiency of ZNF326 induced endothelial senescence, as evidenced by increased SA-β-gal activity and elevated expression of senescence-associated proteins p53 and p21 (Figure 6G-H). Furthermore, ZNF326 knockdown inhibited endothelial cell proliferation and tube formation (Figure S14-15). Taken together, these findings suggest that ZNF326 plays a key role in regulating endothelial cell senescence.

**Figure 6.**
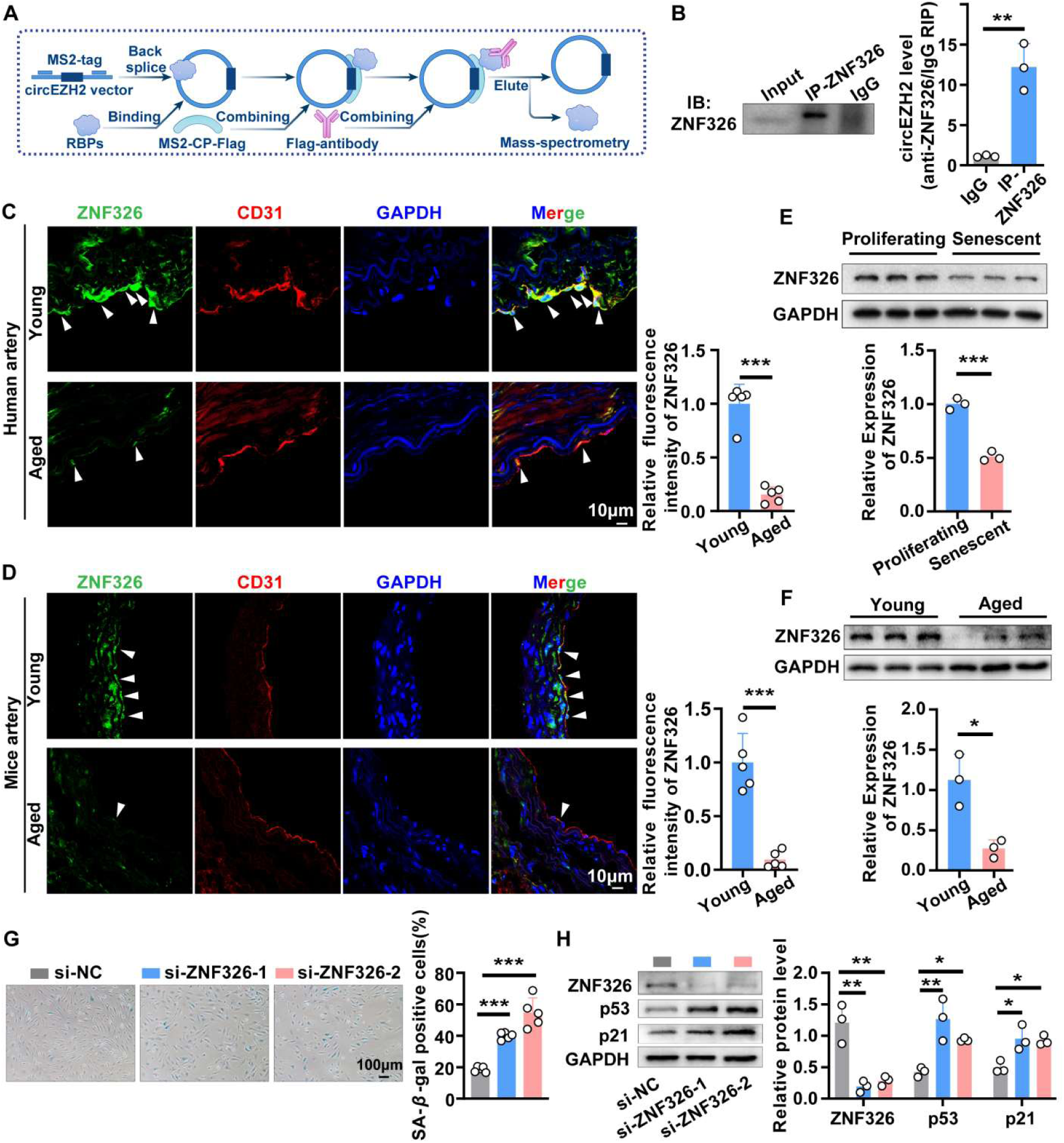
ZNF326 serves as a critical interacting partner of circEZH2 and regulates endothelial cell senescence. **A**, Flow chart shows circEZH2 pull-down assay using the MS2-tagging system. **B**, RNA immunoprecipitation (RIP) assay detecting the interaction between circEZH2 and ZNF326. Cell lysates were incubated with anti-ZNF326 antibody, and bound RNA was extracted for RT-qPCR analysis. **C**, Representative immunofluorescence staining images of CD31 (red), ZNF326 (green), and nuclei (blue) in young and aged human arteries. **D**, Representative immunofluorescence staining images of CD31 (red), ZNF326 (green), and nuclei (blue) in the aortas of young (2 months) and aged (26 months) mice. **E**, Western blot analysis of ZNF326 expression in proliferating and senescent HUVECs. **F**, Western blot analysis of ZNF326 expression in the aortas of young (2 months) and aged (26 months) mice. **G**, SA-β-gal staining in young HUVECs after transfection with si-NC or si-ZNF326-1/2. **H**, Western blot analysis of p53/p21 in young HUVECs after transfection of si-NC or si-ZNF326-1/2. Data are presented as means ± SD, Student’s *t*-test in B, E and F, one-way ANOVA in G and H, \**P* < 0.05, \*\**P* < 0.01, \*\*\**P* < 0.001.

### circEZH2 stabilizes ZNF326 through USP37-mediated deubiquitination

After confirming the interaction between circEZH2 and ZNF326, we further investigated whether this interaction possesses a biological function. Specifically, modulation of circEZH2 expression had no effect on ZNF326 mRNA levels (Figure 7A-B). However, silencing circEZH2 reduced the protein level of ZNF326 in HUVECs, whereas its overexpression led to an increase (Figure 7D-E). Notably, silencing ZNF326 did not alter the expression of circEZH2 (Figure 7C). Collectively, these findings indicate that circEZH2 regulates ZNF326 protein abundance via a post-transcriptional mechanism. Furthermore, an accelerated rate of ZNF326 protein degradation was observed in endothelial cells treated with circEZH2 siRNA and then exposed to the protein synthesis inhibitor cycloheximide (CHX), suggesting that circEZH2 regulates the stability of the ZNF326 protein (Figure 7F). In mammalian cells, the ubiquitin–proteasome system (UPS) plays a crucial role in the degradation of the majority of proteins.^31^ To determine whether the UPS mediated the influence of circEZH2 on ZNF326 degradation, we treated cells with the proteasome inhibitor MG132. We observed that silencing circEZH2 significantly increased the ubiquitination of ZNF326, indicating that circEZH2 stabilizes ZNF326 by inhibiting its ubiquitination (Figure 7G and Figure S16). Previous proteomic analysis has indicated that Ubiquitin-specific peptidase 37 (USP37) might physically interact with ZNF326 in HEK293 cells.^32^ USP37, a deubiquitinase, has been reported to exert its deubiquitinating activity by removing ubiquitin moieties from target proteins.^33, 34^ Thus, we hypothesized that circEZH2 may recruit USP37 to deubiquitinate ZNF326, thereby enhancing its stability. To confirm this, we first demonstrated the interaction between circEZH2 and USP37 using RIP assays (Figure 7H). Subsequently, an exogenous Co-IP assay confirmed that ZNF326 also binds to USP37 (Figure 7I). We further observed that overexpression of USP37 led to an increased expression of ZNF326 in a dose-dependent manner (Figure 7J). Interestingly, knockdown of circEZH2 reduced the interaction between ZNF326 and USP37 (Figure 7K). In summary, circEZH2 may serve as a scaffold to enhance the binding of USP37 and ZNF326, potentially facilitating the effects of USP37 on ZNF326.

**Figure 7.**
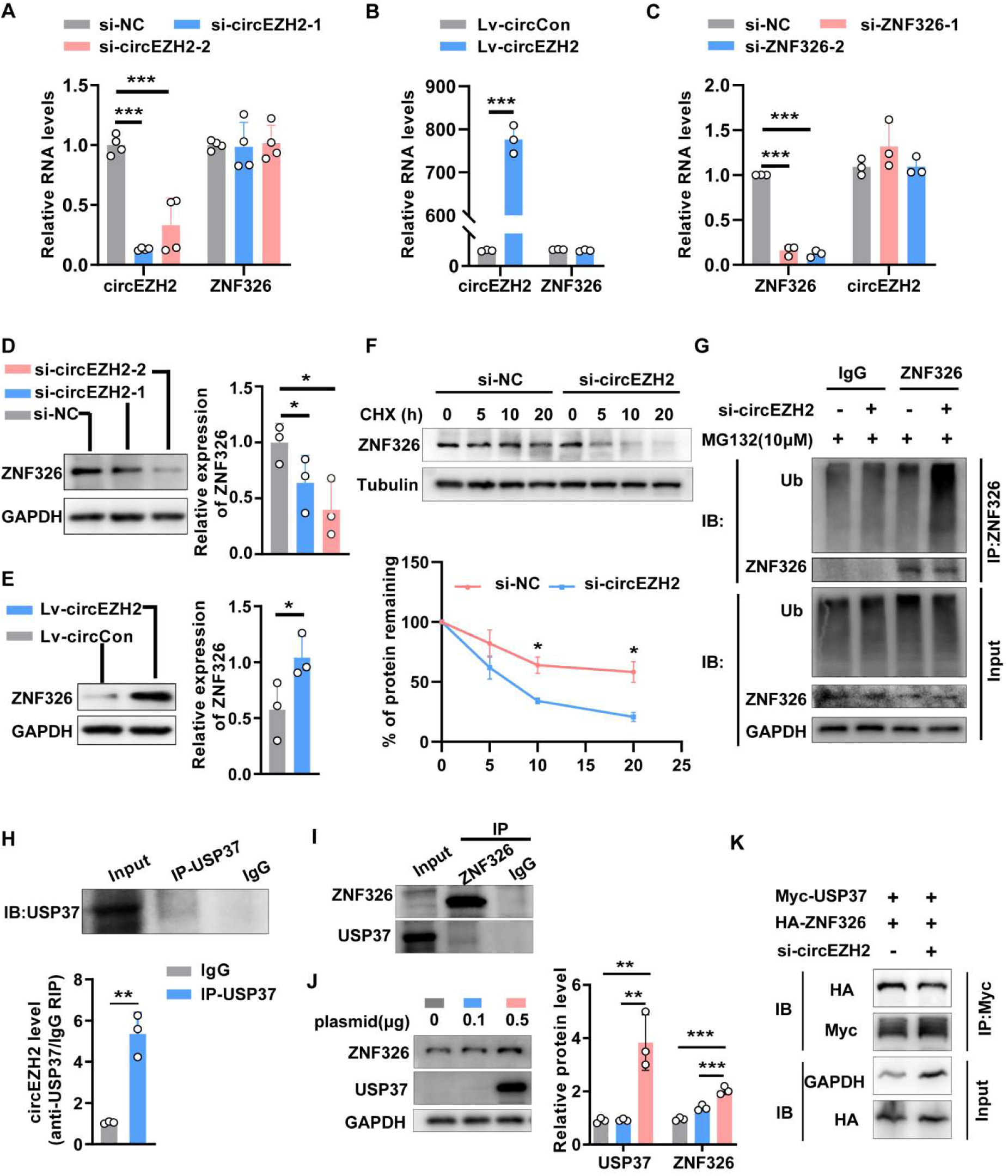
circEZH2 stabilizes ZNF326 through USP37-mediated deubiquitination. **A, B, D, E,** The mRNA and protein levels of ZNF326 were measured by qPCR and Western blot after circEZH2 silencing or overexpression. **C**, The expression of circEZH2 was measured by RT-qPCR after silence of ZNF326. **F**, ZNF326 protein levels at the indicated time points following treatment with cycloheximide (CHX, 10 µg/mL) in transfected HUVECs. **G**, Lysates from circEZH2-knockdown cells were immunoprecipitated (IP) with an anti-ZNF326 antibody and were then used for immunoblot analysis of ubiquitin and ZNF326. **H**, RNA immunoprecipitation detecting the binding of circEZH2 and USP37. Cell lysates were incubated with anti-USP37 antibody, and bound RNA was extracted for RT-qPCR analysis. **I**, Co-IP assays verified the interaction between ZNF326 and USP37 in HUVECs. **J**, Western blot analysis of USP37 and ZNF326 protein levels in HUVECs transfected with increasing doses of USP37 overexpression plasmids. **K**, Co-immunoprecipitation detected the interaction between ZNF326 and USP37 in HEK293T cells transfected with si-NC or si-circEZH2. Data are presented as means ± SD. Student’s *t*-test in B, E, F, H and J; one-way ANOVA in A, C and D. \**P* < 0.05, \*\**P* < 0.01, \*\*\**P* < 0.001.

### Endothelial cell-specific knockdown of ZNF326 counteracts the anti-senescent and anti-atherosclerotic effects mediated by circEZH2 overexpression

To further investigate the role of ZNF326 in the regulation of endothelial cell senescence mediated by circEZH2, we coinjected AAV9-circEZH2 and AAV9-shZNF326 into *ApoE^-/-^*mice. Injection of AAV9-circEZH2 effectively increased the levels of circEZH2 in the intima (Figure S17A). Consistent with the observation *in vitro*, overexpression of circEZH2 also increased the protein level of ZNF326 in the intima, and endothelial cell-specific knockdown of ZNF326 effectively diminished circEZH2-induced ZNF326 upregulation (Figure S17B). Overexpression of circEZH2 reduced SA-β-gal activity in the aortas and aortic roots of *ApoE^-/-^* mice, while also decreasing the protein levels of p53 and p21 in the intimal layer (Figure 8A-F). However, ZNF326 knockdown significantly reversed these effects of circEZH2 overexpression on vascular endothelial cell senescence (Figure 8A-F). We then knocked down ZNF326 in endothelial cells infected with the circEZH2 overexpression lentivirus. Similar to *in vivo* results, overexpression of circEZH2 reduced SA-β-gal-positive cells, enhanced endothelial cell proliferation, and downregulated p53 and p21 levels, while ZNF326 knockdown rescued these effects (Figure 8I-J and Figure S18).

**Figure 8.**
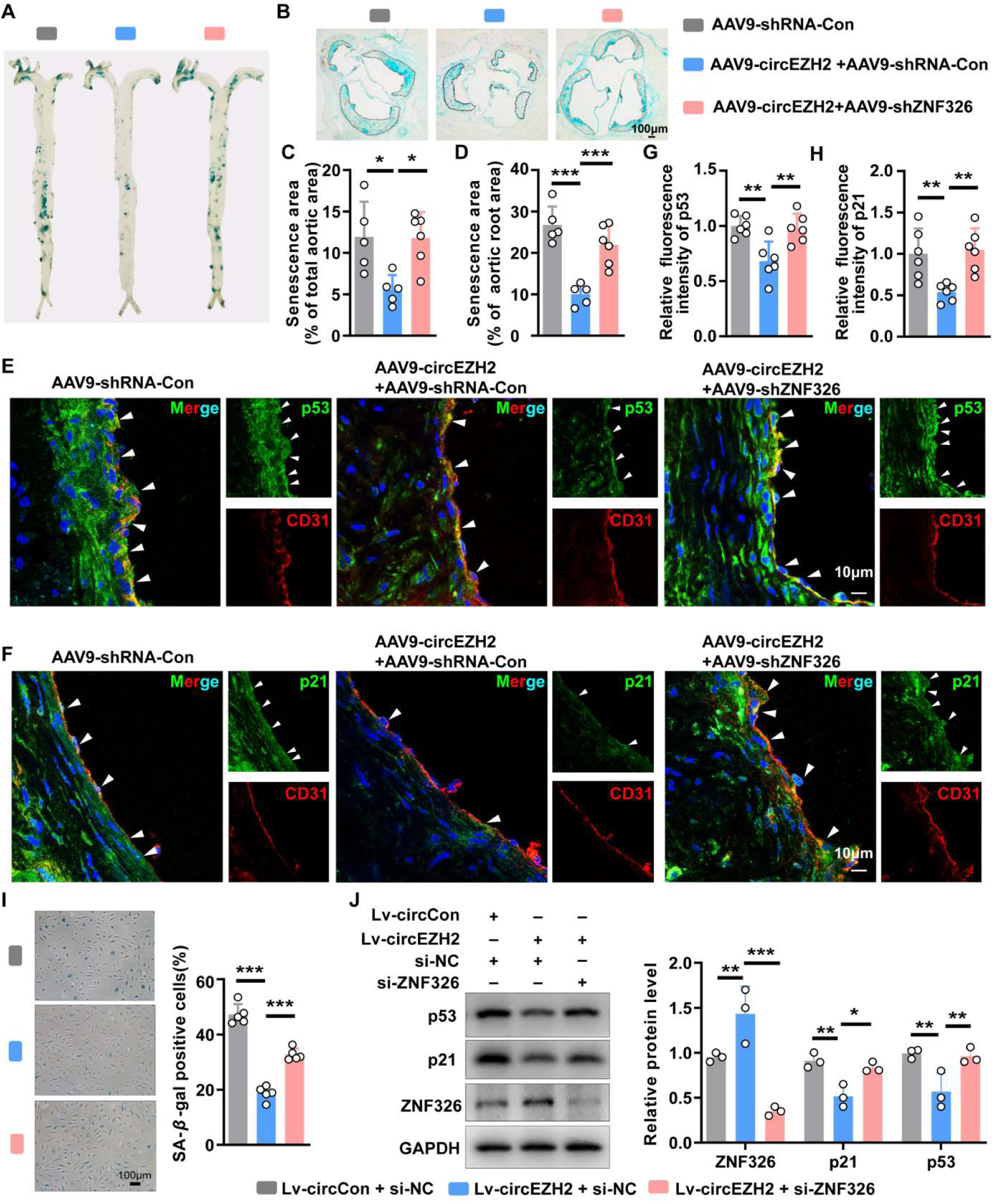
Endothelial ZNF326 knockdown counteracts the anti-senescent effect mediated by circEZH2 overexpression. **A-D**, SA-β-gal staining of whole aortas and aortic roots in *ApoE^-/-^* mice following injection with AAV9-shRNA-Con, AAV9-circEZH2 + AAV9-shRNA-Con, AAV9-circEZH2 + AAV9-shZNF326 (n = 5-6 mice per group). **E-H**, Representative immunofluorescence staining images of CD31 (red), p53/p21 (green), and nuclei (blue) in aortic roots of *ApoE^-/-^* mice from three groups (n = 5-6 mice per group). **I and J**, SA-β-gal staining assays (I), and Western blot analysis of p53/p21 (J) in HUVECs infected with lentivirus expressing circEZH2 followed by knockdown of ZNF326 using siRNA. Data are presented as means ± SD. One-way ANOVA. \**P* < 0.05, \*\**P* < 0.01, \*\*\**P* < 0.001.

In parallel, we assessed the effect of ZNF326 knockdown on the anti-atherosclerotic effects mediated by circEZH2 overexpression. As shown in Figure 9A-C, circEZH2 overexpression significantly reduced the atherosclerotic lesion area compared to the control group. However, when ZNF326 was knocked down in the intima, the lesion area was significantly increased. Upregulation of circEZH2 increased eNOS levels and decreased VCAM1 and ICAM1 levels in the intimal layer, and these effects could be reversed by ZNF326 knockdown (Figure 9D-I). Thus, these results indicate that endothelial cell-specific ZNF326 knockdown abrogates the anti-atherosclerotic effects of circEZH2 overexpression *in vivo*.

**Figure 9.**
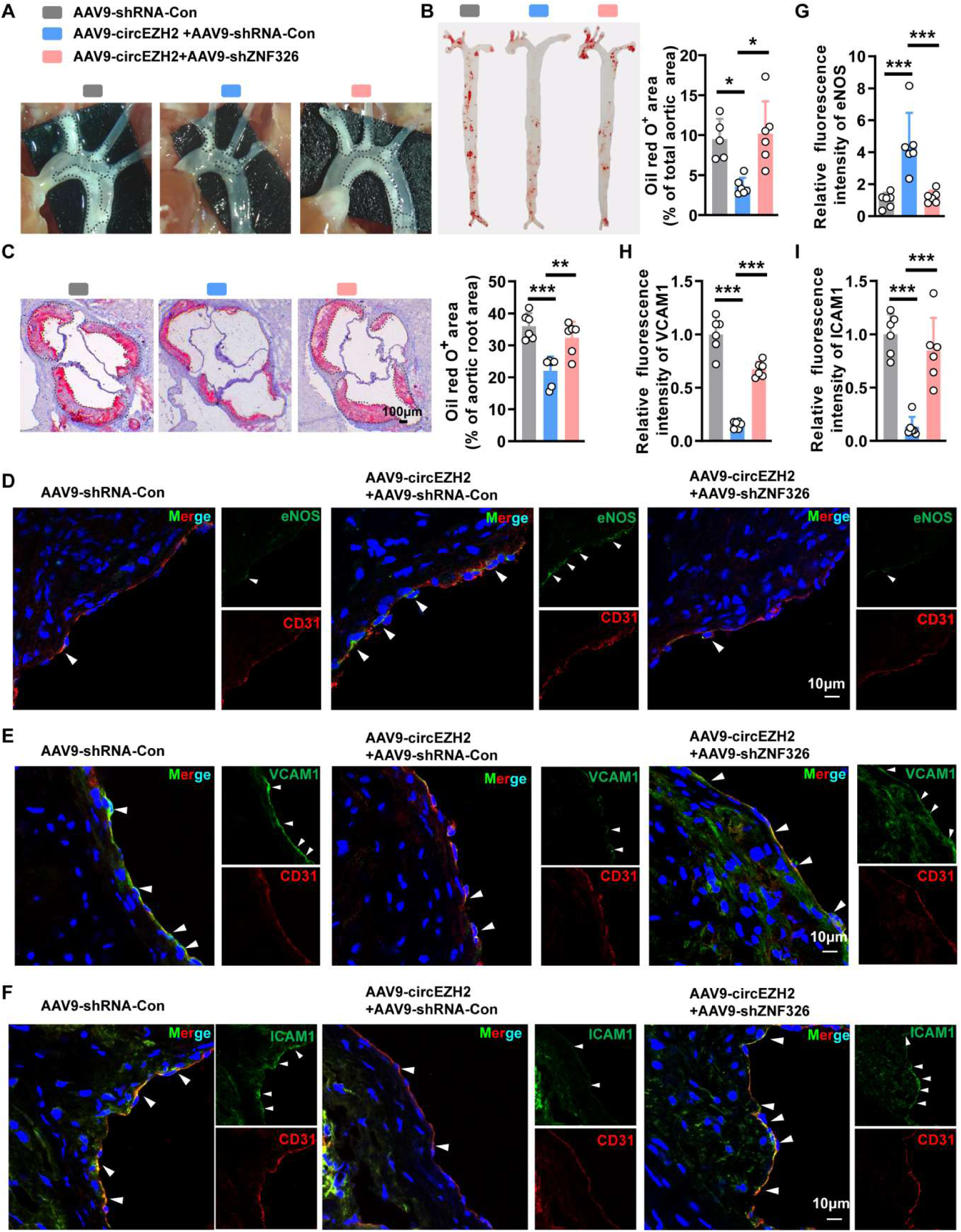
Endothelial cell-specific knockdown of ZNF326 counteracts the anti-atherosclerotic effect mediated by circEZH2 overexpression. **A,** Plaques in the aortic arches of *ApoE^-/-^* mice following injection with AAV9-shRNA-Con, AAV9-circEZH2 + AAV9-shRNA-Con, AAV9-circEZH2 + AAV9-shZNF326. **B and C,** Oil Red O staining of whole aorta and aortic roots of *ApoE^-/-^* mice from three groups (n = 5-6 mice per group). **D-I,** Representative immunofluorescence staining images of CD31 (red), eNOS/VCAM1/ICAM1 (green), and nuclei (blue) in aortic roots of *ApoE^-/-^* mice from three groups (n = 5-6 mice per group). One-way ANOVA. \**P* < 0.05, \*\**P* < 0.01, \*\*\**P* < 0.001.

## Discussion

Vascular endothelial cell senescence induces endothelial dysfunction, which in turn contributes to atherogenesis.^35, 36^ Atherosclerosis is a chronic disorder characterized by fibrofatty accumulations in the walls of arteries, and it is well established as a key etiological factor in a range of cardiovascular diseases.^37^ CircRNAs have emerged as critical regulators in the progression of cardiovascular diseases, and growing evidence implicates their dysregulation in the development of atherosclerosis.^38, 39^ However, the specific role and underlying molecular mechanisms of circRNAs in vascular endothelial cell senescence remain to be fully elucidated. In the current study, we identified a novel endothelial cell senescence-associated circRNA, circEZH2, which undergoes m^6^A modification. The *in vivo* data showed that endothelial cell-specific overexpression of circEZH2 suppresses endothelial cell senescence and atherosclerosis. These findings suggest that circEZH2 plays an important role in endothelial cell senescence and atherosclerosis.

CircEZH2 (hsa_circ_0006357) is a 253-nt circRNA generated by head-to-tail splicing of exons 2 and 3 of the EZH2 gene. It has increasingly been recognized as a functionally diverse circRNA across multiple pathological contexts. A study by Su *et al.* reported that circEZH2 acts as a dual suppressor of miR-363 and miR-708, two microRNAs targeting different regions of EZH2 (3′UTR and coding sequence, respectively), thereby reinforcing EZH2 expression and promoting prostate cancer progression and aggressiveness.^40^ Additionally, circEZH2 sponges miR-133b and prevents IGF2BP2 degradation, which in turn stabilizes m6A-modified CREB1 mRNA and promotes malignancy of colorectal cancer.^41^ Another study has revealed that exosomal circEZH2 alleviates intestinal ischemia/reperfusion injury by interacting with hnRNPA1 and enhancing Gprc5a stability.^25^ Here, we reveal a novel regulatory role of circEZH2 in vascular endothelial cells. We first demonstrate that endothelium-enriched circEZH2 exerts protective effects against endothelial cell senescence and atherosclerotic progression. Our findings expand the repertoire of circEZH2’s regulatory functions from oncogenesis and tissue injury to vascular aging and atherosclerosis.

N^6^-methyladenosine (m^6^A) is one of the most prevalent epitranscriptomic modifications in RNA.^20^ With increasing understanding of m^6^A modification, its regulatory effects on circRNAs have attracted growing attention.^42^ Accumulating evidence indicates that m^6^A-modified circRNAs are involved in diverse biological and pathological processes;^22, 43, 44^ however, their roles in endothelial cell senescence remain unexplored. In the present study, we found that the elevated expression of circEZH2 in young endothelial cells was associated with m^6^A modification and mediated by increased expression of the m^6^A reader IGF2BP2. As a member of the IGF2BP family, IGF2BP2 has been reported to enhance the stability and accumulation of target RNAs by recognizing m^6^A-modified transcripts.^45, 46^ In our data, we demonstrated that circEZH2 was stabilized by IGF2BP2 in a m^6^A-dependent manner.

Endothelial senescence significantly contributes to the development and progression of atherosclerotic plaques,^35^ which has led researchers to focus on developing interventions that can mitigate endothelial senescence to slow plaque growth. In this study, circEZH2 functions as a suppressor of endothelial cell senescence by reducing the expression of cell cycle inhibitors including p53 and p21. We further investigated whether circEZH2 could alleviate atherosclerosis *in vivo*. As anticipated, the overexpression of endothelial circEZH2 not only delayed endothelial cell senescence but also suppressed the progression of atherosclerosis in mice. Notably, endothelial-specific overexpression of circEZH2 did not affect the body weights or lipid profiles of *ApoE^−/−^* mice (Figure S10). Taken together, these results indicate that the anti-atherogenic effects of circEZH2 are likely mediated through the suppression of endothelial cell senescence without affecting lipid metabolism, suggesting that targeting circEZH2 may be a promising strategy for treating atherosclerosis.

CircRNAs have been reported to regulate the expression of their host genes.^47^ However, in this study, circEZH2 did not alter the expression levels of its parental gene, EZH2. These findings indicate that circEZH2 exerts its functions independently of regulating its parental gene in endothelial cells. Growing evidence indicates that certain circRNAs can interact with RNA-binding proteins (RBPs), thereby affecting the function of target protein in diverse physiological and pathological contexts.^48–50^ In the present study, we revealed that circEZH2 primarily exerts its function by binding to ZNF326. ZNF326 is a C2H2-type zinc-finger protein that functions as a key regulator of gene expression.^51^ A study by Yu *et al.* demonstrated that ZNF326 can promote the malignant phenotype of human glioma via the ZNF326-HDAC7-β-catenin signaling pathway.^28^ Rengasamy *et al.* reported that the PRMT5/WDR77 complex regulates alternative splicing and maintains mRNA stability in breast cancer through the symmetric dimethylation of ZNF326.^29^ However, its potential role in endothelial cell senescence or atherosclerosis remains unexplored. In this study, ZNF326 knockdown could mimic the effects of circEZH2 silencing, which promoted endothelial cell senescence and inhibited tube formation. Additionally, endothelial cell-specific knockdown of ZNF326 counteracted the anti-senescent and anti-atherosclerotic effects mediated by circEZH2 overexpression. Collectively, these findings indicate that the circEZH2/ZNF326 axis plays a critical role in regulating endothelial cell senescence and atherosclerosis.

Ubiquitination is one of the most common and important posttranslational modifications of proteins.^52^ Emerging evidence suggests that circRNAs play a critical role in regulating protein stability by affecting the ubiquitination process. For instance, circZBTB46 directly binds to hnRNPA2B1, suppressing its ubiquitination and proteasomal degradation, thereby regulating cell functions and the formation of aortic atherosclerotic plaques.^53^ Additionally, Wang *et al.* reported that circ-0001283 promotes the progression of cardiac hypertrophy by interacting with myosin light chain 3 (MYL3) to inhibit its ubiquitination and enhance its protein expression.^54^ A recent study reported that the long non-coding RNA (lncRNA) OLBC15 enhances the ubiquitin-mediated degradation of ZNF326, thereby promoting the viability, migration, and metastasis of triple-negative breast cancer (TNBC) cells.^55^ To further investigate the molecular mechanism by which circEZH2 knockdown promotes endothelial cell senescence, we explored whether circEZH2 enhances ZNF326 stability by modulating the ubiquitin-proteasome system. Here, we identified that USP37 is a novel binding partner of circEZH2. USP37 is a proteasome-associated deubiquitinating enzyme that protects protein substrates from degradation by removing the ubiquitin chain from substrates bound to the proteasome.^56^ In this study, we found that circEZH2 acts as a scaffold to increase the deubiquitinating enzyme USP37-mediated deubiquitination and stability of ZNF326 protein. Our findings uncover a novel mechanism of ZNF326 regulation and provide strong evidence for the involvement of circRNAs in protein metabolism.

In summary, we identified a novel m^6^A-modified circRNA, circEZH2, that was downregulated in the aged aortic intima and advanced plaques. CircEZH2 was stabilized by m^6^A reader IGF2BP2. Functionally, endothelial cell-specific overexpression of circEZH2 delayed senescence and attenuated atherosclerosis progression. Mechanistically, circEZH2 functions as a scaffold, facilitating the binding of USP37 to ZNF326 and thereby enhancing the stability of ZNF326 protein. Collectively, these findings support the potential of circEZH2-based gene therapy as a novel therapeutic approach for atherosclerosis.

## Acknowledgments

The authors thank X.J., X.M., Z.J., Z.S. and Y.L. for conceptualizing and designing the study, performing most experiments, interpreting the results, and writing the manuscript; Z.L., M.C. and X.T. for collecting clinical samples; Y.Q., Z.X., Y.X., Y.-F.Q, S.W., L.M., H.L. Z.W. and X. L. for providing constructive suggestions on experimental design and revising the manuscript; and X.X. and J.T. for directing the study, providing scientific input and data analysis, and preparing the manuscript. All authors read and commented on the article.

## Sources of Funding

This work was supported by the National Natural Science Foundation of China (82571787, 82071576, 82270460), Guangdong Basic and Applied Basic Research Foundation (2026A1515010440), Guangdong Province Innovative Team Project for Ordinary Universities (2021KCXTD049), the Medical Scientific Research Foundation of Guangdong Province of China (A2024346), the Discipline Construction Project of Guangdong Medical University (4SG24018G, 4SG22306P), Special Project for Clinical and Basic Sci&Tech Innovation of Guangdong Medical University (GDMUZDCG25001, GDMULCJC2025121, GDMULCJC2024097), and National College Students’ Innovation and Entrepreneurship Training Program (202410571001).

## Disclosures

The authors declare no conflict of interest.

## Supplemental Material

Supplemental Methods

Table S1-S13

Figures S1–S18

Major Resources Table

ARRIVE Guidelines

## Nonstandard Abbreviations and Acronyms

circRNA: circular RNA
MeRIP: methylated RNA immunoprecipitation
m6A: N6-methyladenosine
IGF2BP2: insulin-like growth factor 2 mRNA-binding protein 2
USP37: ubiquitin-specific peptidase 37
ZNF326: zinc finger protein 326
eNOS: endothelial nitric oxide synthase
ICAM1: intercellular adhesion molecule 1
VCAM1: vascular cell adhesion molecule 1
RBP: RNA-binding protein
HAEC: human aortic endothelial cell
HCAEC: human coronary artery endothelial cell
HUVEC: human umbilical vein endothelial cell
ApoE: apolipoprotein E
CHX: cycloheximide
Co-IP: coimmunoprecipitation
AAV9: adeno-associated virus serotype 9
HFD: high-fat diet
RIP: RNA immunoprecipitation
RNA-FISH: RNA fluorescence in situ hybridization
RT-qPCR: reverse transcription quantitative polymerase chain reaction
SA-β-gal: senescence-associated β-galactosidase

## Supporting Information

Supporting Information is available from the Wiley Online Library or from the author.

## Notes

### Competing Interest Statement

The authors have declared no competing interest.

